# Reduced PDE4D expression and activity in Acrodysostosis Type 2 patient fibroblasts underlie disease pathology

**DOI:** 10.64898/2026.08.10.743905

**Authors:** Oliver FW Gardner, Jiayue Ling, Tawan Munkongcharoen, Elka Kyurkchieva, Harry G Leitch, Louise C Wilson, George S Ballie, Patrizia Ferretti

## Abstract

**Background:** Acrodysostosis type 2 (ACRDYS2) is a rare autosomal dominant disease characterized by skeletal defects and cognitive deficit, with clinical symptoms observed in multiple other tissues including the skin. It is caused by mutations in a phosphodiesterase, PDE4D, a key regulator of cAMP/PKA (cyclic adenosine monophosphate / protein kinase A) signalling. Despite its well-defined genetic causes, the molecular mechanisms underlying the disease remain poorly understood, with studies based largely on engineered cellular models reaching conflicting interpretations.

**Methods:** To **investigate** how endogenous dynamics are affected by PDE4D mutations in unmanipulated cells, we studied PDE4D transcript and protein expression, activity and downstream signalling in native dermal fibroblast from ACRDYS2 patients and healthy controls.

**Results:** Significant reduction in total PDE4D expression in patient cells was observed both at the transcript and protein level, with **marked** decreases in the long isoforms PDE4D4 and PDE4D7; a reduction in *PDE4D9* mRNA was also observed. PDE4D enzymatic activity was reduced in ACRDYS2 fibroblasts, though total PDE activity was largely preserved. Reduced PDE4D expression was associated with an increase in the phosphorylated form of the cAMP-responsive transcription factor CREB and elevated *PRKAR1A* (PKA type 1 regulatory subunit alpha) transcript levels, suggesting altered downstream signalling. Interestingly, expression of the related phosphodiesterase family member PDE4B was increased, consistent with a compensatory response to reduced PDE4D function.

**Conclusions:** This is the first study demonstrating reduced PDE4D expression and isoform-specific dysregulation in native ACRDYS2 cells. Together, our results support a model in which reduction in PDE4D activity and compensatory changes in other PDE4 family members contribute to the molecular pathology of ACRDYS2, providing new insights into the molecular mechanisms underlying this disorder.

## Introduction

Acrodysostosis (ACRDYS) is a very rare autosomal dominant disease belonging to a larger group of disorders known as iPPSDs (inactivating PTH/PTHrP signalling disorders). Mutations in two genes, *PRKAR1A* (type-1A regulatory subunit of protein-kinase-A, PKA) and *PDE4D* (cAMP-specific-phosphodiesterase-4D), cause two forms of the disease, ACRDYS1 (also known as IPPSD4) and ACRDYS2 (also known as IPPSD5) respectively [1–4]. Skeletal defects, particularly in facial bones (e.g. midface hypoplasia), hands and feet (brachydactyly), and vertebral column (e.g. spinal stenosis) as well as increased obesity, are observed in both ACRDYS types. However, other features are more commonly observed in either ACRDYS1 or 2. For example, resistance to hormonal stimulation in the growth plate of developing bones and short stature are more commonly present in ACRDYS1, whereas cognitive deficits are more frequently apparent in ACRDYS2. It should also be noted that several other tissues, including the skin, are often affected in ACRDYS patients, as reported in a survey conducted by the Acrodysostosis Patient Charity, but they have yet to be studied [5, 6].

Both PRKAR1A and PDE4D are key components of the cAMP/PKA signalling pathway which is involved in the response to certain hormones (e.g. parathyroid hormone) via activation of G-protein-coupled receptors. In both forms of ACRDYS gene mutations have been thought to lead to reduced cAMP/PKA signalling and ultimately a reduction in CREB activation and downstream gene expression. In ACRDYS1, *PRKAR1A* mutations result in the failure of this regulatory subunit to be activated by cAMP and release the PKA catalytic subunit, leading to reduced cAMP/PKA signalling [7]. In contrast, the mechanism(s) by which mutations in PDE4D lead to a reduction in cAMP/PKA signalling have been more controversial [1]. Regulation of PDE4D is complex, with fine control of cAMP microdomains depending on phosphorylation, dimerization and compartmentalization of the enzyme. Further complexity is added by the 11 *PDE4D* splice variants that contain either both UCR1 (upstream conserved region 1) and UCR2 (upstream conserved region 2) domains (long form), the UCR2 domain only (short form) or a truncated UCR2 domain (super short form); these functional domains together with the catalytic domain are also hot spots for *PDE4D* mutations reported in ACRDYS2 patients [1, 8, 9].

Several different molecular mechanisms of the diseases have been suggested. Gain of function has been suggested by analysis of PDE4D activity following transfection of CHO cells with PDE4D3, one of the long PDE4D isoforms, carrying mutations in different domains [10]. However, in the only ACRDYS2 study on patient cells, where EBV-immortalized lymphocytes were used, PDE4D mutations seemed to result in reduced enzymatic activity, which would be expected to increase cAMP availability and consequently increase signalling. The net reduction in activity in ACRDYS2 was suggested to be due to overcompensation by other PDE4 isoforms [11]. It has also been suggested that mutations associated with acrodysostosis affect removal of cAMP from PRKAR1A, hence prolonging PKA activation, though PDE8 rather than PDE4D was investigated in this study [12].

Together most models used so far to study human ACRDYS2, such as overexpression of mutated PDE4Ds, are unlikely to precisely represent the level of nuanced regulation that cells have for controlling cAMP, and this could explain some of the discrepancy in the literature. The use of patients’ cell carrying PDE4D mutations has been hindered by the rarity of the disease. Availability of such cells is crucial to addressing how mutations affect expression of PDE4 isoforms, enzymatic activity, dimerization of longform PDE4Ds, potential compensatory mechanisms, as well as possible changes in intracellular localization, which is crucial to fine regulation of cAMP signalling.

We have been able to access skin biopsies from consenting ACRDYS2 patients and generated three dermal fibroblast lines from patients with different mutations, that for the first time have allowed in parallel characterisation of non-manipulated multiple lines. Importantly, this has allowed us to address the gain versus loss of function hypothesis in cells from a tissue that can be affected in patients. We demonstrate for the first time in ACRDYS2 patients’ dermal fibroblasts significant reduction at the transcriptional and protein level of certain PDE4D isoforms, reduced enzymatic activity, and possibly some compensation from other PDE4 family members.

## Methods

### Fibroblast isolation and culture

Three ACRDYS2 patient fibroblast cell lines were studied. One was kindly provided by the Rare & Inherited Disease Laboratory of the North Thames Genomic Laboratory Hub (A-F1), and the other two (A-F2 and A-F3) from surplus skin tissue from consenting patients under ethical approval from the Camden and Islington Community Local Research Ethics Committee (London, UK). Clinical genetic diagnoses showed that patients carried the following mutations: A-F1 carried a PDE4D c.935T>C; p.(Leu312Pro) mutation; A-F2 a PDE4D c.1772C>A p.(Thr591Asn) and A-F3 a PDE4D c.934C>T p.(Leu312Phe) mutation. Mutations in patient fibroblast lines generated were confirmed by Sanger sequencing. Three human dermal fibroblast cell lines were used as healthy controls (C-F1, C-F2, C-F3). They were derived from skin biopsies from foetuses at 13, 17 and 18 post conception weeks of development provided by the Human Developmental Biology Resource (HDBR) under ethical approval, respectively. In some experiments also a fibroblast line from a consented paediatric patient undergoing surgery and established by Prof. Wei-Li Di was used (F48).

To isolate fibroblasts from skin biopsies, samples were first washed in DMEM Glutamax (Gibco) with 100 µg/ml Primocin (InvivoGen). The tissue was then minced with a scalpel before digestion in 0.5% Trypsin-EDTA (Gibco) for one hour at 37°C. The tissue was then washed with DMEM Glutamax with Primocin and 10% Fetal Bovine Serum Value (FBS, Gibco) and then digested for ninety minutes in DMEM Glutamax with 1.5 mg/ml Collagenase from Clostridium histolyticum (Sigma-Aldrich). The resulting cell suspension was washed and then resuspended in DMEM Glutamax with Primocin and 10% FBS and then plated in a T25 Flask (Corning). Cells were left for forty-eight to seventy-two hours to attach before media was changed to DMEM Glutamax with 100 U/ml Penicillin-Streptomycin (Gibco) and 10% FBS (proliferation medium). Media was changed three times a week and when fibroblasts reached confluence they were passaged using 0.05% trypsin-EDTA (Gibco). Following isolation lines were cultured in proliferation media and tested for mycoplasma contamination using the MycoAlert™ PLUS Mycoplasma Detection Kit (Lonza) according to the manufacturers’ instructions, before freezing in 90% FBS with 10% DMSO (ChemCruz) and storage in liquid nitrogen.

For experimental work fibroblasts were thawed into proliferation medium at a density of 3,000 cells/cm^2^ and expanded for one to two passages before use. For immunocytochemistry fibroblasts were plated at 3,000 cells/well into 8-well chambered culture slides (Falcon) and allowed to proliferate for 48-96 hours. For cAMP quantification and PDE activity measured using the cAMP-Glo™ Assay (Promega) CF-1 and AF-1 fibroblasts were seeded at 10,000 cells/well in a µCLEAR® white CELLSTAR® 96 well tissue culture treated plate (Greiner bio-one) and harvested for analysis 24 hours later. For gene expression analysis, Western blotting and PDE4 activity measurement all lines control and patient lines were thawed and expanded before seeding at 2,000-3,000 cells/cm^2^ into 6-well plates (Corning) for gene expression analysis and 10cm tissue culture dishes for Western blotting and PDE4 activity measurement. Fibroblasts in 6-well plates and 10cm dishes were then expanded in parallel to 95-100% confluence before harvesting.

### RNA isolation, reverse transcription and quantitative real time PCR

Wells of six-well plates containing fibroblasts from all control and patient fibroblast lines grown for gene expression analysis were lysed with 1 ml of Trizol (Invitrogen) which was then collected and stored at -80°C before further processing. Five samples were collected for CF-1, CF-2, CF-3, AF-1 and AF-2, and six collected for AF-3. Samples were thawed and chloroform (Fisher Scientific) was added to induce phase separation. Following shaking and centrifugation the aqueous phase was collected and RNA precipitated using an equal volume of isopropanol (Acros Organics). The RNA pellet was then washed two times with 75% ethanol (Sigma-Aldrich) and resuspended in RNAse free water (Severn Biotech). RNA quantity and purity were determined using a NanoDrop™ One spectrophotometer, 1 µg of RNA from each sample was then treated with DNAse I (Thermo Fisher Scientific) according to manufacturer’s instructions. Reverse transcription was then performed using random hexamer primers and the RevertAid First Strand cDNA Synthesis Kit (Thermo Fisher Scientific) according to the manufacturer’s protocol and the cDNA was then stored for future use at -80°C.

Quantitative real time PCR (qPCR) was performed using the QuantiTect™ SYBR® Green PCR Kit (Qiagen), to each well 5 µl of QuantiTect™ SYBR® Green was mixed with 3.3µl of RNAse free water and 0.7 µl of 10 µM primer mix and 1 µl of cDNA. The primers listed in Table 1 were used to detect the expression of either all of the members of each PDE4 subfamily (Pan-PDE4A, Pan-PDE4B, Pan-PDE4C or Pan-PDE4D) or individual long isoforms of PDE4D (PDE4D3, PDE4D4, PDE4D5, PDE4D7, PDE4D8, PDE4D9) as well as protein kinase regulatory subunit 1-alpha (PRKAR1A), Sox9 and RPL19A. Primers were designed and tested at the UCL GOSH ICH and synthesised by Sigma-Aldrich. Samples were analysed using the 7500 Fast Real Time PCR System (Applied Biosysyems). Data were analysed using the ΔΔCt method, the ΔCt for all samples was calculated by normalising to the expression of RPL19A and the ΔΔCt calculated by normalising to the average ΔCt of all control samples. Fold-change was then calculated using the formula 2^ΔΔCt^.

**Table 1.**
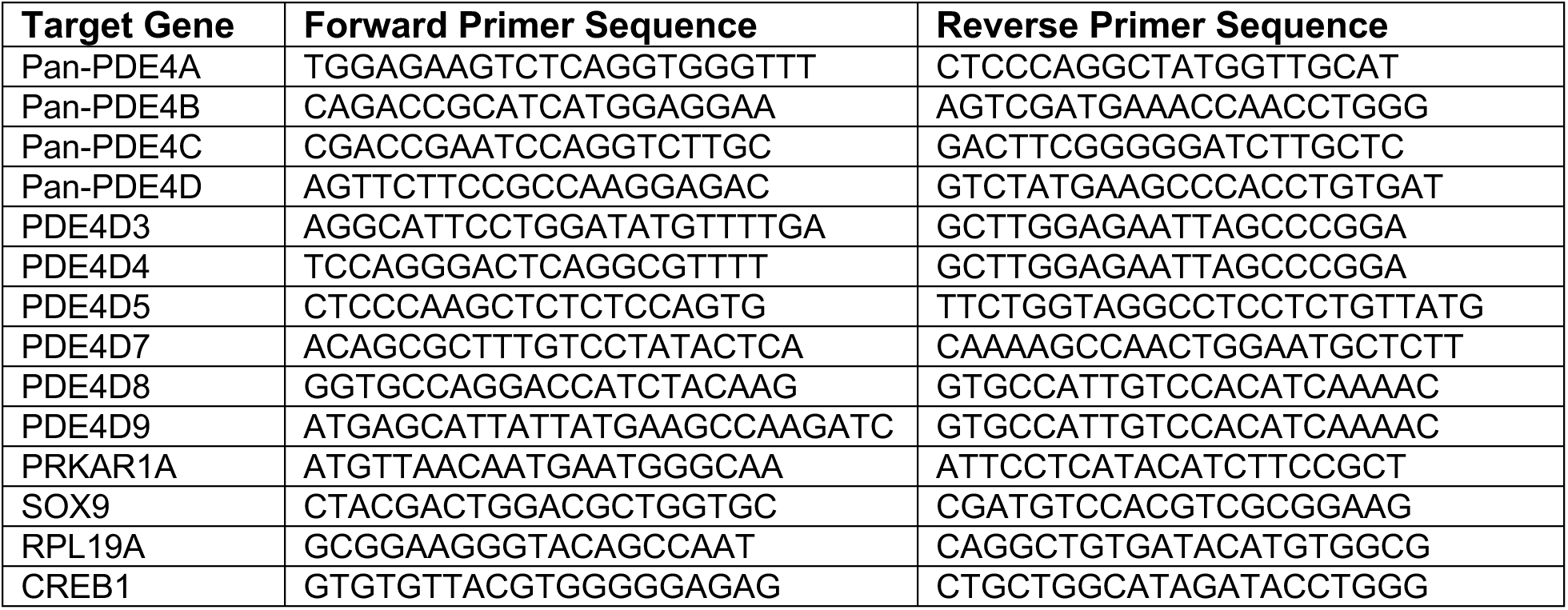
Primer sequences used for gene expression analysis by reverse transcription quantitative PCR.

### Immunocytochemistry

Fibroblasts grown in chamber slides for immunocytochemistry (ICC, both n=3) were washed twice with Dulbecco’s phosphate buffered saline (DPBS, Gibco) before fixation in 10% neutral buffered formalin (Sigma-Aldrich) for ten minutes. This was followed by three washes in phosphate buffered saline (Oxoid), before permeabilization 0.1% Triton-X100 in PBS for seven minutes. Wells were then washed twice with PBS before blocking for one hour in PBS with 10% FBS and 2% bovine serum albumin (Sigma Aldrich). Slides were then washed before incubation with the primary antibody, Pan-PDE4D (1:200; Proteintech) and Phalloidin 594 (1:400; Invitrogen) diluted in PBS with 5% FBS and 1% BSA for 1 hour. After washing with PBS, slides were incubated with the secondary antibody (Donkey anti-Rabbit 488 (Invitrogen) in PBS with 5% FBS and 1% BSA for 1 hour, then washed and incubated with Hoeschst dye (10 mg/mL, Thermo Fisher) in PBS with 5% FBS and 1% BSA for ten minutes. After washing, slide were mounted using Vectashield Antifade Mounting medium (2B scientific). Negative controls where the primary antibody was omitted were included in each experiments (Supplementary Figure 1). A Zeiss Observer fluorescence microscope with a Hamamatsu Flash 4v3 camera was used to image fluorescently labelled sections at 63x magnification and images processed using FiJi software [13].

### Western Blotting

Fibroblasts from all lines were expanded in 10 cm dishes before harvesting for Western blotting, when they reached 95-100% confluent the proliferation medium was removed and the plates were washed with DPBS. All the DPBS was then removed and the plates were frozen dry at -80 °C where they were stored until further processing.

**Table 2.** Primary and secondary antibodies used for western blotting.

| Antibody name | Host species | Supplier | Dilution |
| --- | --- | --- | --- |
| **Primary** |  |  |  |
| PDE4B (Pan) | Goat | In-house | 1:5000 |
| PDE4D (Pan) | Goat | In-house | 1:5000 |
| PDE4D7 | Goat | In-house | 1:1000 |
| PDE7B (Pan) | Rabbit | Abcam (ab170914) | 1:5000 |
| PDE8A (Pan) | Rabbit | Proteintech (13956-01-AP) | 1:1000 |
| Phospho-CREB | Rabbit | Proteintech (28792-1-AP) | 1:500 |
| Total CREB | Mouse | Proteintech (67927-1-Ig) | 1:1000 |
| Phospho- (Ser/Thr)<br>PKA substrate | Rabbit | Cell Signaling Technology<br>(9621) | 1:1000 |
| PKA RβII | Mouse | Thermo Fisher, #PA5-13793 | 1:1000 |
| PKA Cβ | Rabbit | BD Transduction (610626) | 1:1000 |
| GAPDH | mouse | Proteintech (60004-1-Ig) | 1:40,000 |
| **Secondary** |  |  |  |
| Donkey anti-rabbit 800 | Donkey | LICORbio (926-32213) | 1:10,000 |
| Donkey anti-mouse 800 | Donkey | LICORbio (926-32212) | 1:10,000 |
| Goat anti-rabbit 680 | Goat | LICORbio (926-68071) | 1:10,000 |
| Donkey anti-mouse 680 | Donkey | LICORbio (926-68072) | 1:10,000 |
| Donkey anti- goat 800 | Donkey | Abcam (ab186699) | 1:10,000 |
| Donkey anti-goat 680 | Donkey | Abcam (ab175776) | 1:10,000 |

Cells were lysed in 3T3 lysis buffer (25 mM HEPES, 50 mM NaCl, 50 mM NaF, 30 mM Na4P2O7, 5 mM EDTA, 10% glycerol, 1% Triton, pH=7.5) supplemented with cOmplete EDTA-free Protease inhibitor Cocktail tablet (Roche) and PhosSTOP tablets (Roche). Protein concentration was determined by Bradford protein assay. Protein samples were prepared in SDS buffer and denatured at 95°C for 5 minutes; 40µg of protein were loaded into 4-12% SDS-PAGE Bis-Tris Gels (ThermoFisher Scientific), gels were run at 150V for 10 minutes and 200V for 50 minutes with NuPAGE MOPS SDS Running Buffer (Invitrogen). Proteins were transferred from the gels to nitrocellulose membranes (ThermoFisher Scientific) at 25V for 90 minutes in NuPAGE Transfer Buffer (Invitrogen). Membranes were blocked in 0.5x Intercept® blocking buffer (LI-COR) for 1 h at room temperature. Antibodies were prepared in 0.5x Intercept® antibody diluent. The primary and secondary antibodies used for western blotting are listed in Table 2. Membranes were incubated with the primary antibody overnight at 4°C, while the secondary antibody was added for 1 h at room temperature. Membranes were washed three times for 5 min each in 1xTBST (Tris-Buffered Saline with Tween 20) after each incubation with antibodies. Membranes were visualized via Odyssey CLx Imager (LI-COR) and analysed using Image Studio software.

### PDE4 Activity Measurement (Radio assay)

For PDE activity measurement, fibroblasts from all lines were prepared and stored as described for western blotting. PDE activity was indirectly measured through direct quantification of radioactively tagged 8-[^3^H] adenosine from cAMP hydrolysis by PDE [14].

Cells were lysed in KHEM buffer (50 mM KCl, 50 mM HEPES, 10 mM EGTA and 1.9 mM MgCl2 + 1x cOmplete EDTA-free Mini Protease Inhibitor Cocktail tablet; Roche Diagnostics) and lysed by going through 10 rounds of freeze on dry ice followed by thawed at 37°C cycle. The Bradford assay was used to determine the protein concentration of each sample. Protein samples were diluted to a final concentration of 80 μg in Buffer A (20 mM Tris–HCl, pH 7.4). Samples were prepared in triplicate to a final volume of 50 μL, either in the absence or presence of rolipram, a selective phosphodiesterase-4 (PDE4) inhibitor (10 μM). This concentration exceeds the reported IC_50_ for PDE4 isoforms (∼2 μM) (Soderling et al., 1998) and was selected to ensure near-complete inhibition of PDE4 activity. Furthermore, 10 μM rolipram has been widely used in biochemical PDE assays to achieve maximal PDE4 inhibition (Xin et al., 2015), The assay was initiated by adding a cAMP substrate mixture consisting of 3 μL of radiolabelled 8-[^3^H]-cAMP (“hot”) and 2 μL of 1 mM unlabelled cAMP (“cold”) to each sample. Samples were incubated for 10 minutes at 30 °C, after which the reaction was terminated by boiling for 2 minutes, followed by cooling on ice for 15 minutes. Snake venom was then added to a final concentration of 1 mg mL^−1^, and samples were incubated for a further 10 min at 30 °C. Subsequently, 400 μL of Dowex anion exchange resin slurry (resin_2_O, 1:1:1) was added to each sample. Samples were vortexed and incubated on ice for 15 min, with mixing every 5 minutes, before centrifugation at 20,210 × g for 3 min at room temperature. A 150 μL aliquot of the supernatant was transferred to 1 mL of Opti-Flow SAFE 1 scintillation cocktail. Additional control samples containing 50 μL of the cAMP substrate mixture were prepared to determine the total counts per minute (CPM). Radioactivity was quantified using a Tri-Carb 4810TR Liquid Scintillation Counter (Revvity, MA, USA) to determine the amount of hydrolysed 8-[^3^H]-cAMP.

PDE activity was calculated as pmol cAMP hydrolysed min^−1^ mg protein^−1^ using the following equation:

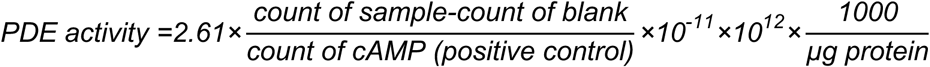

where 2.61 corrects for the fraction of supernatant recovered following anion exchange chromatography, 10^−11^ accounts for the incubation time and the amount of cAMP substrate used, 10^12^ converts moles to picomoles, and the factor (1000/\mu\text) normalizes activity to milligrams of protein. PDE4-specific activity was determined by subtracting the PDE activity measured in the presence of rolipram from the total PDE activity measured in its absence.

### cAMP Quantification

The total cAMP content of C-F1 (n=6) and A-F1 fibroblasts (n=6) was determined using the cAMP-Glo™ Assay (Promega). Measurements were made according to the manufacturer’s instructions, except that duplicate assays were performed firstly in the presence of the broad spectrum PDE inhibitor 3-Isobutyl-1-methylxanthine (IBMX, 100 µM) and the PDE4 inhibitor rolipram (500µM) and secondly in the presence of IBMX alone without rolipram. Measurements were made using a FLUOstar OPTIMA plate reader (BMG Labtech).

### Statistical Analysis

Statistical analysis was performed GraphPad Prism 9 software (GraphPad Software Inc.). For gene expression results comparing control fibroblast lines (n=3) against patient lines (n=3) normality was tested using the D’Agostino-Pearson test. Where samples were normally distributed (Pan-PDE4A, B, C, D, PDE4D3 and 5) significance was tested using an unpaired 2-way T-Test. Where distribution was not normal (PDE4D9, PRKAR1A and Sox9) significance was tested using an unpaired 2-tail Mann-Whitney test.

For gene expression results comparing individual fibroblast lines (n=5 or 6 samples per line) normality was tested using the Shapiro-Wilk test. Where samples were normally distributed (Pan-PDE4A and C) significance was tested using one-way ANOVA with Tukey’s multiple comparison test. Where distribution was not normal (Pan-PDE4B, D, PDE4D3, 5, 9, PRKAR1A and Sox9) significance was tested using the Kruskal-Wallis test with Dunn’s multiple comparison test. Gene expression signal for PDE4D4 and PDE4D7 was detected in all control samples but in too few of the patient samples for statistical analysis. PDE4D8 was excluded from statistical analysis as no signal was detected in any control or patient samples.

For western blotting, the desired band intensity was normalised to loading control protein GAPDH; unpaired t test was performed between control and patient group. For PDE activity assay, after the activity was calculated, unpaired t test was performed between control and patient group. Results of cAMP quantification in CF-1 and AF-1 (n=6 for both) were tested for significance using an unpaired 2-way T-Test.

Differences were considered significant when p< 0.05.

## Results

Mutations in the ACRDYS2 fibroblasts studied were confirmed by sequencing and a summary of the mutations and clinical presentation for each patient is shown in Table 3. All patients presented brachydactyly and facial dysplasia; intellectual disability was reported in the two patients aged eight and twenty-one but could not be assessed in the patient who died aged four months.

**Table 3.** Summary of PDE4D mutations and clinical presentation in ACRDYS2 patients from whom fibroblasts were generated from skin biopsies.

| ACRDYS2 Fibroblasts | Age * | Sex | Mutation | Affected region | Clinical presentation |
| --- | --- | --- | --- | --- | --- |
| A-F1 | <1 | M | c.935T>C;<br>p.(Leu312Pro) | UCR2 | Antenatal oligohydramnios, intrauterine growth retardation, brachydactyly, nasal and maxillary hypoplasia, extensive lymphangiectasis of intra-parenchymal and pleural lymphatic vessels. Died age 4 months from acute pneumonia a pulmonary hypertension |
| A-F2 | 8 | F | c.1772C>A;<br>p.(Thr591Asn) | Catalytic | Generalised marked brachydactyly, marked nasal hypoplasia, mild mid-face hypoplasia, pale irides, moderate intellectual disability, obstructive sleep apnoea, otitis media with effusions and conductive hearing impairment, hypermetropia, idiopathic intracranial hypertension requiring VP shunt, chronic constipation, raised PTH but normal calcium and thyroid function |
| AF-3 | 21 | M | c.934C>T;<br>p.(Leu312Phe) | UCR2 | Generalised marked brachydactyly, marked nasal hypoplasia, moderate mid-face hypoplasia, pale irides, moderate intellectual disability with autism and ADHD, non-verbal, epilepsy, undescended testes, obstructive sleep apnoea, otitis media with effusions and conductive hearing impairment, pancreatitis secondary to gallstones, normal PTH & thyroid axis to date |

\*: years at time of biopsy

### Total PDE4D and long isoform expression in healthy and ACRDYS2 fibroblasts

The relative gene expression levels of the PDE4D and its long isoforms were determined in three control and three patient fibroblast lines using either primers detecting either all of the PDE4D isoforms (‘pan’) or specific long isoforms. Analysis of total *PDE4D* expression by RT-qPCR showed consistently lower transcript levels (P<0.0001) in all fibroblast lines derived from patients carrying *PDE4D* mutations as compared to controls (Figure 1). Given the existence of several long isoforms of PDE4D, we investigated whether any variant was particularly affected in patients’ cells with a focus on the long forms, which regulate activity via dimerization and phosphorylation of UCR1 by PKA. Within the long isoforms of *PDE4D* no significant difference was detected in the expression of *PDE4D3* (P=0.5346) or *PDE4D5* (P=0.9076) between control and patient lines. *PDE4D9* expression was significantly lower in patients’ fibroblasts than controls (P<0.0001). For *PDE4D4* and *PDE4D7* there was an almost total absence of expression in patient samples (transcript detected in only 1 of 16 samples and 3 of 16 samples, respectively, across three cell lines), despite detectable expression in control samples from all three control lines (10 of 15 samples and 14 of 15 samples respectively). This indicates a clear reduction in the expression of *PDE4D4* and *PDE4D7* in patient cells, which could not be tested for statistical significance for this technical reason. No signal was detected for *PDE4D8* in either control or patient lines, suggesting it is not highly expressed in fibroblasts.

**Figure 1.**
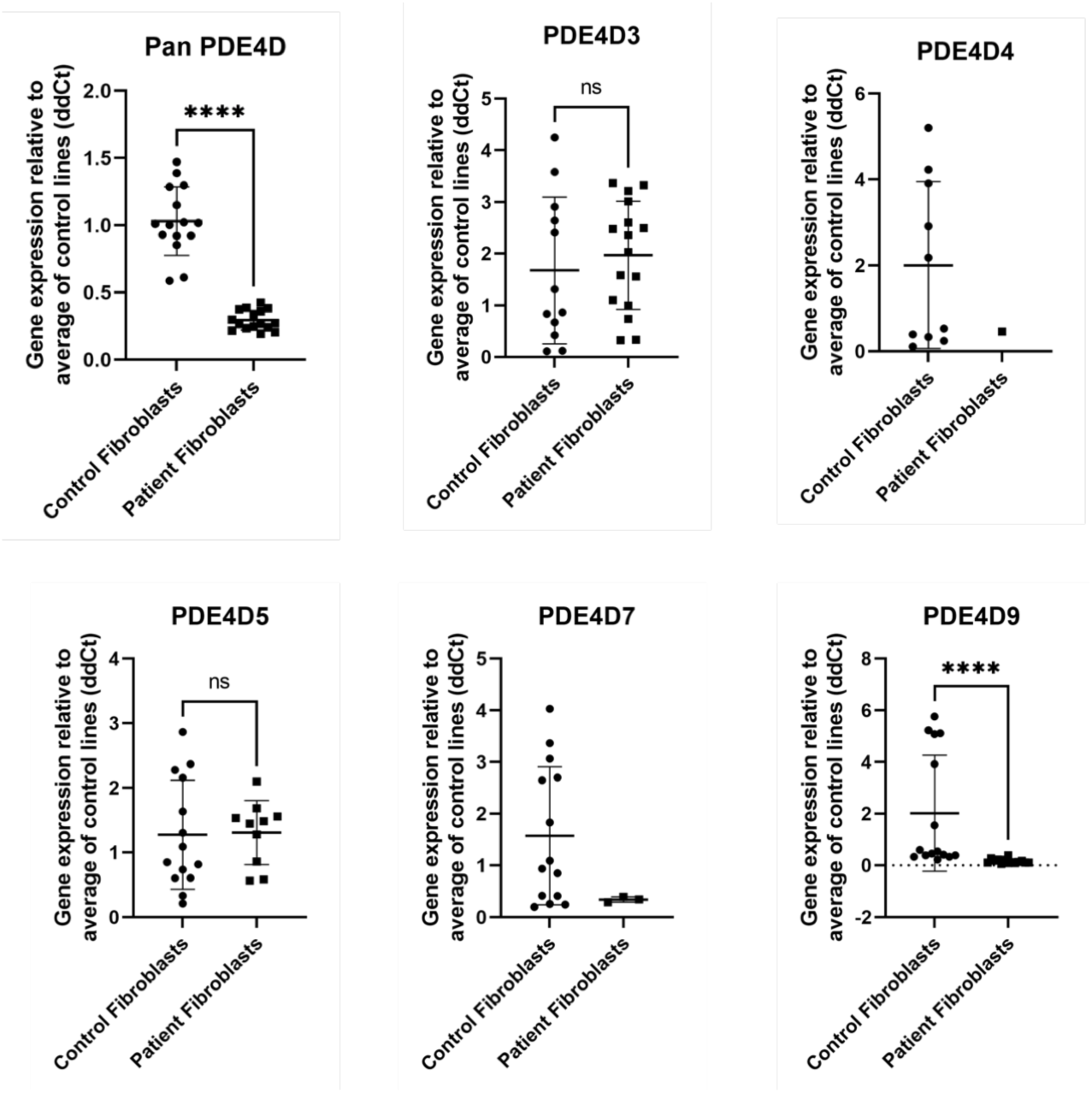
Changes in PDE4D isoform transcript expression in healthy and ACRDYS2 fibroblasts. Total PDE4D transcript levels are significantly reduced in ACRDYS fibroblasts. PDE4D4 is detectable at low levels only in 1 of 16 ACRDYS2 samples and PDE4D7 only in 3 out of 16. PDE4D9 is also significantly reduced.

We then assessed whether these changes in PDE4D gene expression were reflected by changes at the protein level by western blotting (Figure 2). Consistent with the transcript data, a significant decrease in PDE4D in ACRDYS2 fibroblasts was detected using a pan-PDE4D antibody Figure 2A). Furthermore, the PDE4D7 variant was also greatly reduced in patients’ cell as compared to healthy fibroblasts Figure 2B). In contrast no significant change was seen in PDE4D5, which was hardly detectable (not shown).

**Figure 2.**
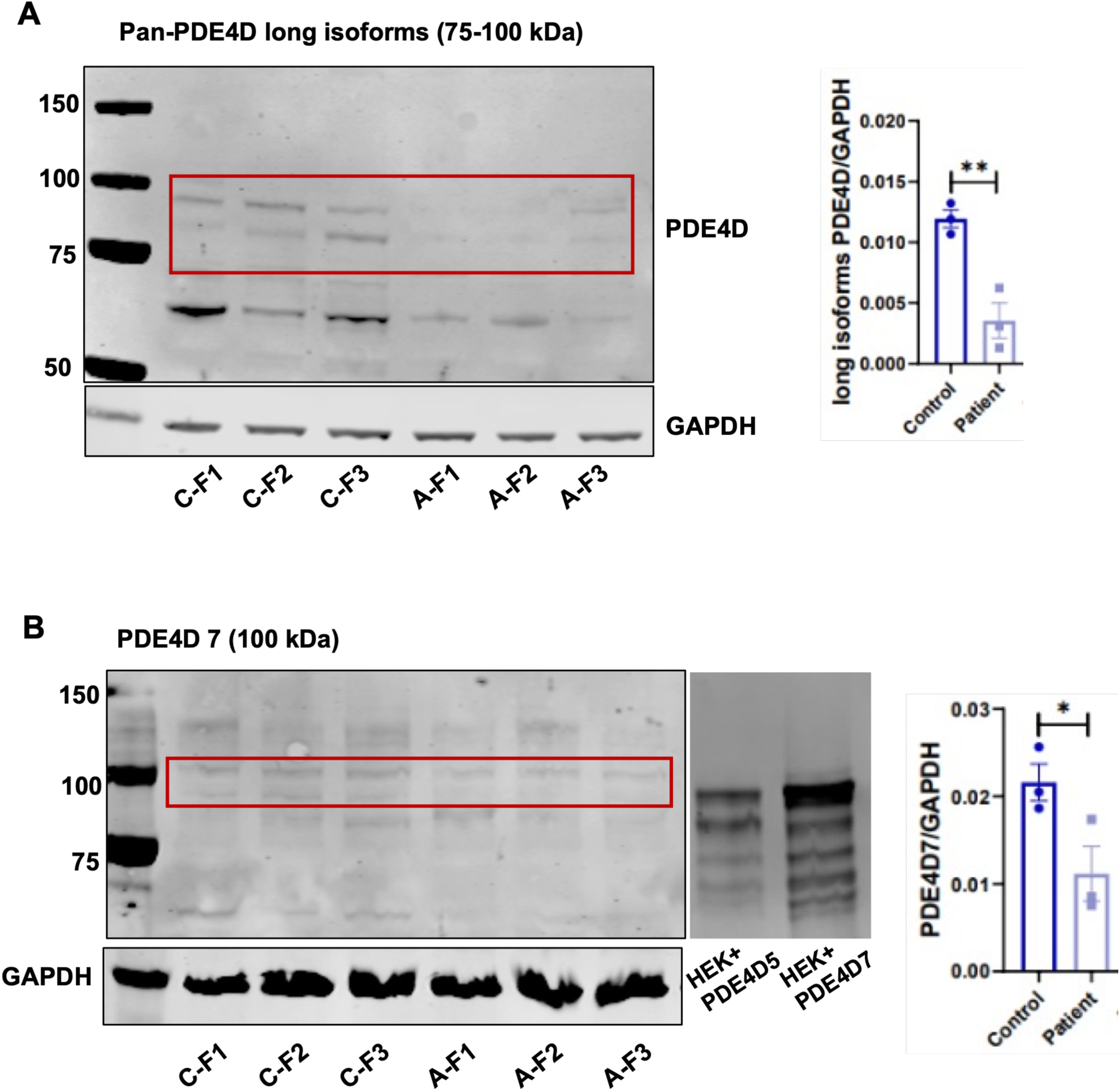
Changes in PDE4D protein in healthy and ACRDYS2 fibroblasts assessed by western blot. **A)** Total PDE4D protein levels are significantly reduced in ACRDYS fibroblasts as indicated by densitometric analysis with PDE4D normalized to GAPDH. **B)** PDE4D7 protein is also significantly reduced. *: p<0.05; **: p<0.01.

Finally, we immunostained healthy and ACRDYS2 fibroblasts with an antibody to PDE4D to assess whether there was any obvious change in PDE4D intracellular localization in diseased cells (Figure 3). Variation in staining intensity among cells was observed in both healthy and ACRDYS2 fibroblasts, with some clusters of strongly positive cells observed also in some ACRDYS2 fibroblasts, particularly in A-F1. However, no evidence of changes in PDE4D intracellular compartmentalization in ACRDYS2 fibroblasts was apparent.

**Figure 3.**
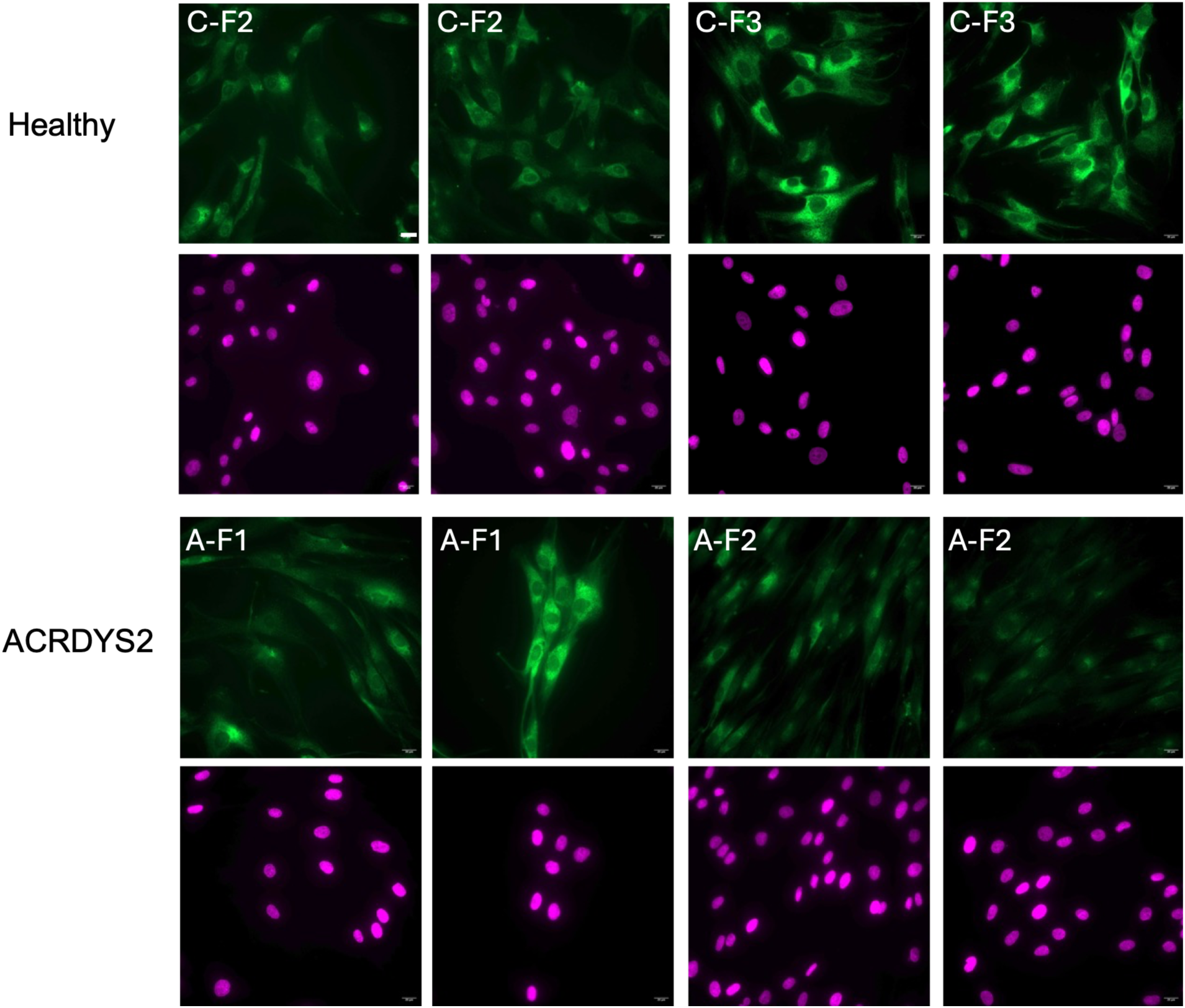
PDE4D immunostaining in healthy and ACRDYS2 fibroblasts. Images exemplify differences in stain intensity observed both in healthy (C) and ACRDYS2 (A) fibroblasts within the same culture. PDE4D is in green and nuclei are in magenta. Scale bars= 20 µm.

### Changes in PDE4D activity in healthy and ACRDYS2 fibroblasts

We then wished to establish whether PDE and PDE4 activity were affected in ACRDYS2 fibroblasts. The extent of cAMP hydrolysis was assessed in control and ACRDYS2 cells by a radioassay assessing PDE and PDE4 activity (Figure 4A). Whereas total PDE activity was similar in control and ACRDYS2 fibroblasts, PDE4 specific activity appeared to be reduced in ACRDYS2, as indicated by the much lower levels of cAMP hydrolized in two (A-F1 and A-F2) of the three patients’ lines as compared to controls. This finding was further validated by measuring the levels of cAMP in F1 and A-F1 fibroblasts using the cAMP-Glo™ Assay (Figure 4B). The assay was performed both with the dual inhibitors IBMX, a non-selective PDE inhibitor, and Rolipram, a PDE4 inhibitor, and with only IBMX. No significant difference was detected between samples measured with IBMX and rolipram (P=0.0638) together, suggesting similar overall levels of intracellular cAMP in the control and patient sample. However, when the assay was performed in the absence of the PDE4 inhibitor rolipram there was an increase in the luminescence in both control and patient cells which reflects a decrease in cAMP as a result of the activity of PDE4. The reduction in cAMP levels was significantly greater in control than in the patient cells (P<0.0001), suggesting reduced PDE4 activity in the patient fibroblasts.

**Figure 4.**
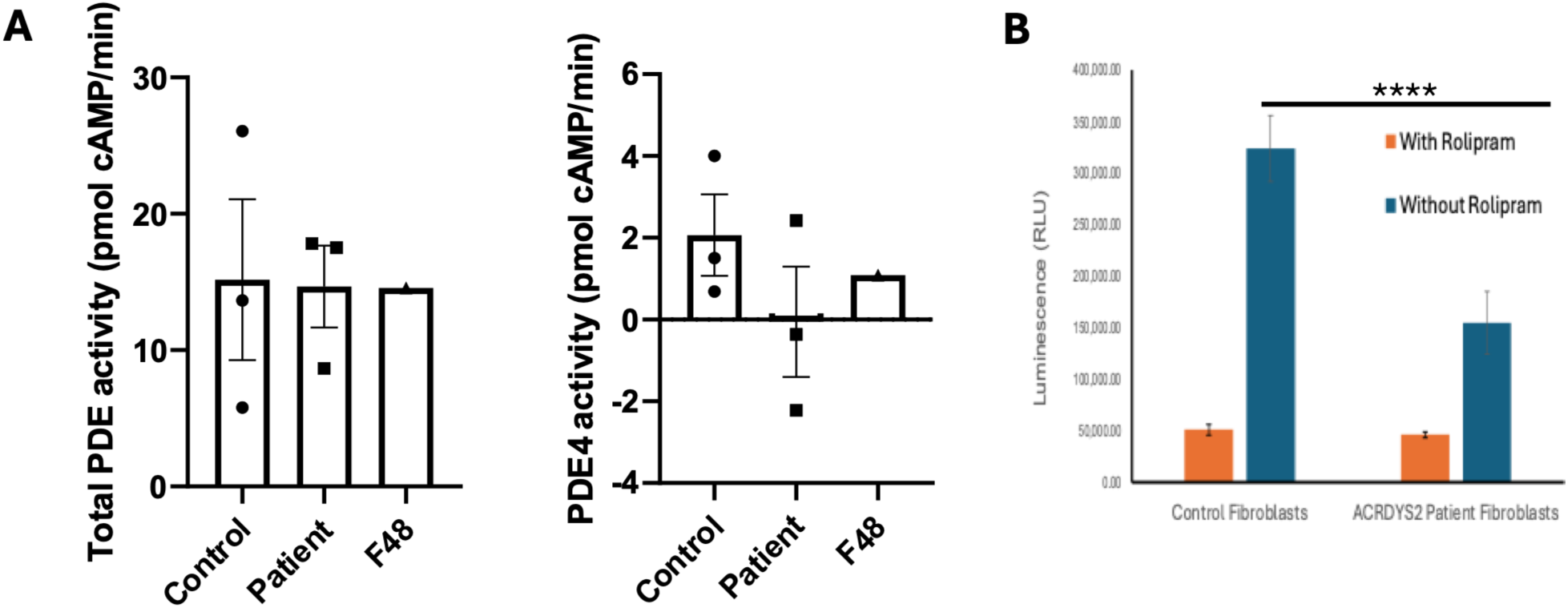
Changes in PDE and PDE4 activity and in cAMP levels in control and ACRDYS2 fibroblasts. A-B). PDE and PDE4 activity measurement by a radio assay in control (C-F1, 2, 3 and F48) and ACRDYS2 (A-F1, 2, 3) fibroblasts. PDE activity is comparable in control and patient fibroblasts whereas PDE4 activity is greatly reduced. **C)** Total cAMP content in C-F1 and A-F1 fibroblasts (n=6). Increased luminescence in both control and patient cells without rolipram treatment is indicative of decreased cAMP levels. Note that cAMP levels are significantly higher in patient than control fibroblasts (****p<0.0001).

As it has been suggested that the net result of PDE4D mutations in ACRDYS2 patients is a reduction in cAMP signalling, but we had found PDE4D to be downregulated and less active (suggesting increased cAMP signalling), we carried out some functional assays. Analysis of CREB phosphorylation was assessed in all ACRDYS2 and healthy lines C-F1,2,3, and in an additional line from a patient with an unrelated disease. As shown in Figure 5A-B, although the ratio between pCREB and CREB was not affected in the ACRDYS2 lines, the level of pCREB was significantly higher in patient than in control fibroblasts, suggesting increased downstream signalling as well as increased CREB levels. Analysis of *CREB1* transcript did not show any difference in gene expression between control and ACRDYS2 fibroblasts (Figure 5C).

**Figure 5.**
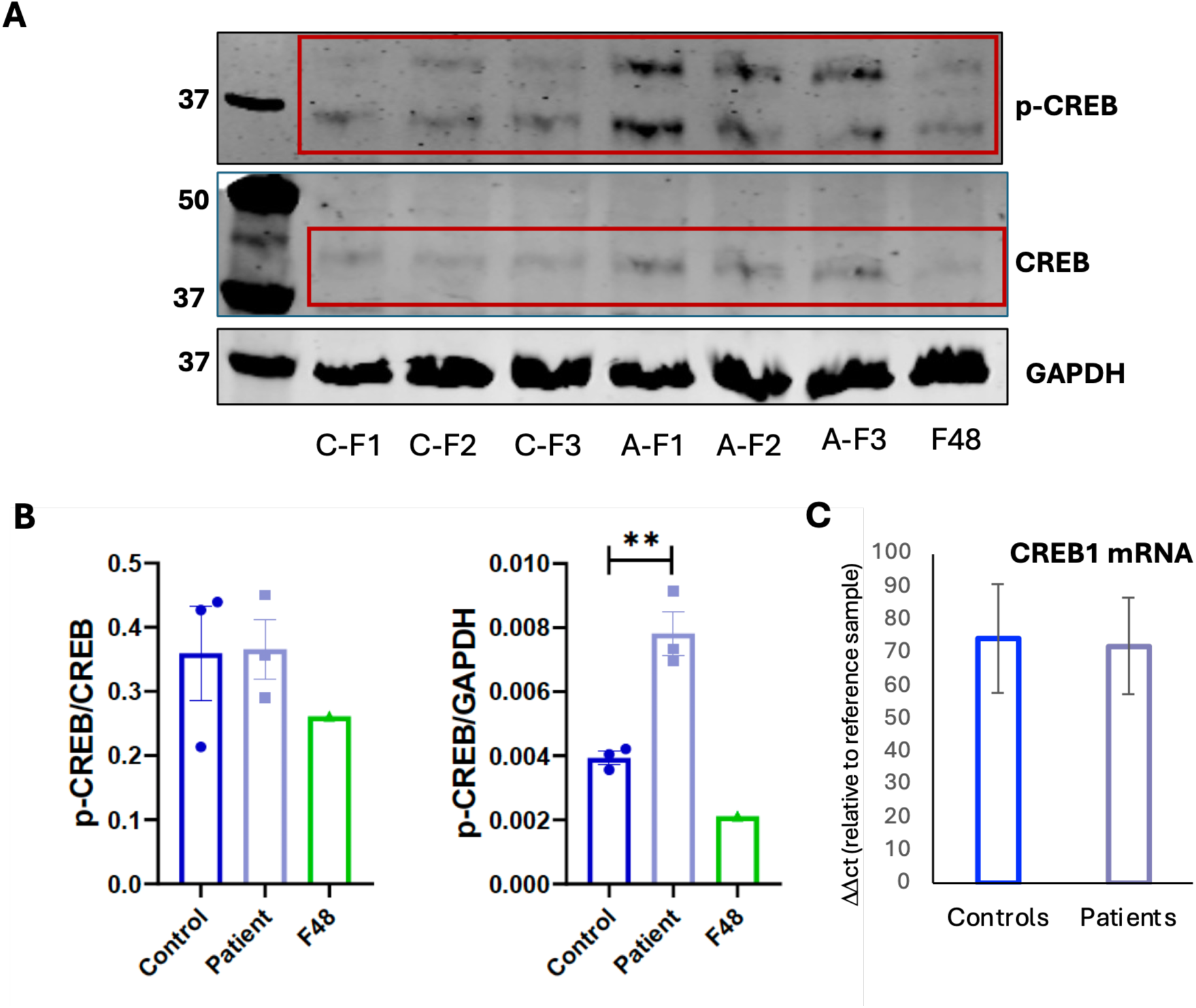
Analysis of CREB and CREB phosphorylation (p-CREB) in control and ACRDYS2 fibroblasts. **A)** Western blot of control (C-F1, 2, 3 and F48) and patient (A-F1, 2, 3) cells for CREB and p-CREB. **B**) Densitometric analysis of western blots normalized to GAPDH; note comparable pCREB/CREB ratio in control and patient cells but increased pCREB in ACRDYS2 fibroblasts. **C)** Analysis of CREB1 mRNA by RTqPCR; no significant difference in transcript level between lines is observed. **: p<0.01.

As SOX9 is a CREB responsive transcription factor also involved in skeletal development, we assessed its expression in healthy and ACRDYS2 fibroblasts (Figure 6A). SOX9 transcript levels were lower in patient lines (M ± SD: control 1.841 ± 1.50 and patient 0.684 ± 0.37), though their reduction did not reach statistical significance (P=0.1627), possibly due to large variability. We also assessed expression of the *PRKAR1A* transcript (Figure 6B), which inhibits the PKA catalytic subunit until it binds cAMP, and found it to be significantly increased in patient lines (P=0.0003). In contrast, western blot analysis showed no significant difference in the regulatory, PKA-RβII, and catalytic, PKA Cβ, subunits of PKA at the protein level, though a trends toward reduction in ACRDYS2 fibroblasts was apparent (Figure 6C-D). Also some reduction in phosphorylated PKA (p-PKA) substrates was observed in ACRDYS fibroblasts, but it was not statistically significant (Figure 7).

**Figure 6.**
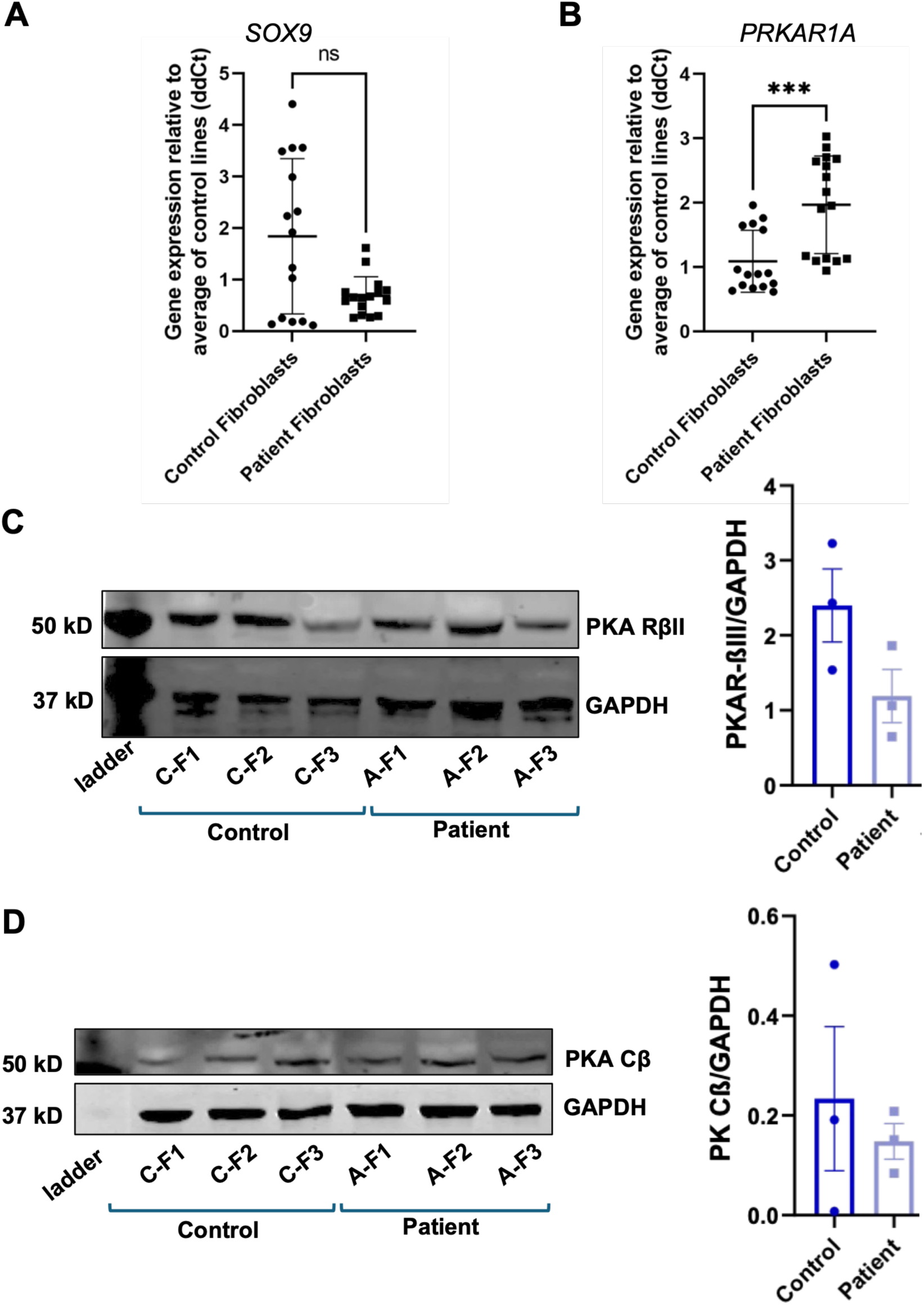
Changes in SOX9 and PRKAR1A expression in healthy and ACRDYS2 fibroblasts. A-B) Detection of SOX9 and PRKAR1A transcripts by RT-qPCR: a significant increase in PRKAR1A transcript levels is observed in ACRDYS fibroblasts (***: p=0.0003). **C-D)** Western blot and densitometric analysis of PKA-RßII and PKA Cß normalized to GAPDH. No significant difference in expression is observed between control and ACRDYS2 fibroblasts.

**Figure 7.**
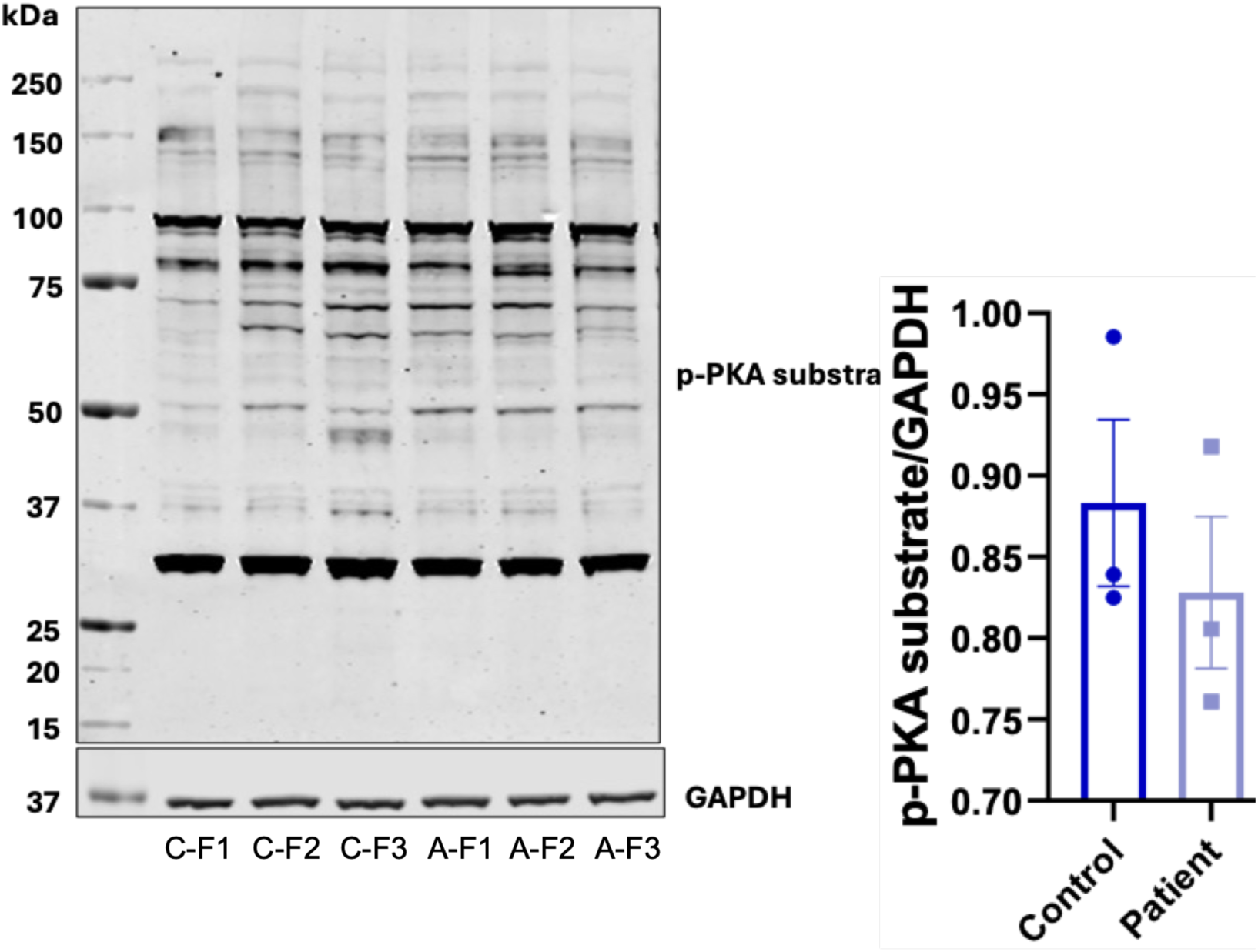
Detection of proteins phosphorylated by PKA in healthy and ACRDYS2 fibroblasts. No statistically significant change in staining of the antibody to phosphorylated PKA (p-PKA) substrate is observed in ACRDYS fibroblasts, as indicated by densitometric analysis normalized to GAPDH, though the trend is toward reduced phosphorylation.

Finally, to establish whether the reduction in PDE4D might activate compensatory mechanisms, we investigated expression of the other PDE4 subfamily members, PDE4A, B, and C in the three control and three patient fibroblast lines using primers detecting all members of each PDE4 subfamily (Figure 8A). No significant transcriptional change was detected in the expression of PDE4A between control and patient lines (P=0.1245) and PDE4C was reduced in ACRDYS2 lines (P=0.0184).). In contrast, notwithstanding high variability, PDE4B expression was significantly elevated in patient lines (P=0.0179). A significant increase in PDE4B expression in ACRDYS2 was also observed at the protein level as shown by western blot (Figure 8B).

**Figure 8.**
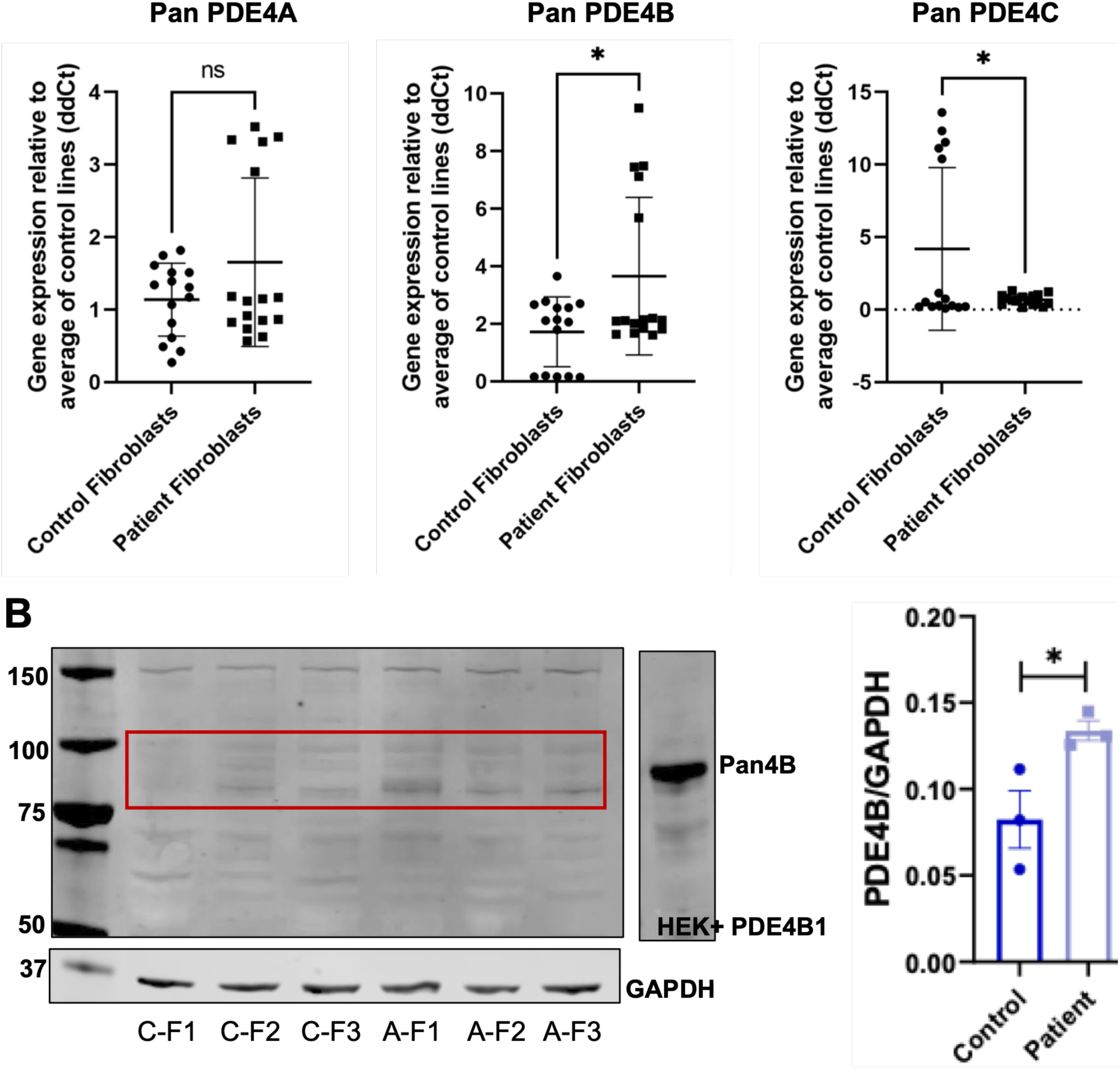
PDE4 subfamily members expression in healthy and ACRDYS2 fibroblasts. **A)** Expression of PDE4A, B, and C assessed by RT-qPCR in control (C-F1, 2, 3) and patient (A-F1, 2, 3) fibroblast lines. PDE4B is significantly increased, whereas PDE4C is decreased in patient cells, notwithstanding variability among lines. **B)** Expression of PDE4B in the same 3 control and 3 ACRDYS2 cell lines assessed by western blot. Densitometric analysis of western blot normalized to GAPDH shows a significant increase in PDE4B in patient cells. *: p<0.05.

## Discussion

This is the first study investigating the effects of PDE4D mutations in multiple ACRDYS2 and healthy lines without transfections or other manipulations, hence allowing us to assess endogenous dynamics in patient cells. It has shown that PDE4D mutations in fibroblast from three patients lead to a reduction in both PDE4D gene and protein expression, which had not been previously reported, independent of the mutation that they carry.

Furthermore, this study has shown that the main PDE4D isoforms affected in ACRDYS2 are PDE4D4 and PDE4D7, with PDE4D9 also showing some reduction in its expression in patient fibroblasts. In contrast, RNA expression of other long isoforms was not significantly different, though there was some variability among ACRDYS2 lines which might be either due to differences in genetic background or different mutations, or a combination of the two.

No obvious change in PDE4D intracellular localization was observed in ACRDYS2 cells, suggesting that PDE4D compartmentalization is not affected by the mutations found in our patients. However, the immunostaining pattern was heterogeneous, with a subset of cells showing stronger immunoreactivity than others in both control and patient-derived fibroblasts. This may partly be explained by the presence of other bands on western blots suggesting that the antibody may recognize additional protein isoforms. Furthermore, variability in staining intensity may be linked to differences in the cell physiological state, such as proliferation or migration [15, 16].

Analysis of intracellular localization will deserve further investigation, particularly because, in other disorders caused by PDE mutations, disease pathogenesis has been linked to PDE10A mislocalization leading to enhanced protein degradation [17].

### PDE4D transcription is reduced in ACRDYS2 fibroblasts whereas PDE4B is increased

The reduction in *PDE4D* transcription was surprising and the mechanisms leading to it have yet to be elucidated. Activated PKA is known to directly phosphorylate the PDE4D UCR1 domain to boost its catalytic activity and rapidly clear cAMP [18], but it is not known to directly affect *PDE4D* transcription. On the other hand, *PDE4D* expression is regulated by multiple promoters, some containing CREB binding sites [19, 20]. Although *PDE4D4* expression levels are not believed to be regulated by cAMP, as treatment with dibutyryl-cAMP upregulated transcription of only *PDE4B2* and short *PDE4D* isoforms, an increase in cAMP has been found to reduce *PDE4A10* transcription in cardiomyocytes, and CREB has been reported to regulate a *PDE4B* intronic promoter [21, 22]. This is relevant to our study, and consistent with the activation of a compensatory feedback mechanism in which enhanced cAMP/CREB signalling drives *PDE4B* transcription to limit cAMP accumulation in patient fibroblasts. It is also tempting to speculate that the increase in phosphorylated CREB we have observed in ACRDYS2 fibroblasts might lead to recruitment of transcriptional regulators or DNA modifiers at the *PDE4D* promoter, or of nearby histones leading to PDE4D down-regulation. It has been suggested that methylation at the *PDE4D* promoter decreases *PDE4D* expression in airways smooth muscle cells [23]. Interestingly, acetylation as well as methylation of histones close to the *PDE4D* gene promoter region has been reported to reduce PDE4D expression [24].

Whichever mechanism might lead to transcriptional regulation of *PDE4D* and *PDE4B* in patient cells, the concordance in changes in PDE4D and PDE4D7 expression at the transcriptional and protein level suggests that protein clearance/turnover is not affected by the ACRDYS2 mutations, and that regulation of expression levels occurs predominantly at the transcriptional level. These findings are important because reduction in PDE4D was not previously considered as a factor underlying the diseases [25].

### PDE4D protein level and signalling are affected by ACRDYS2 mutations

We have shown that both PDE4D protein expression and overall PDE4 activity are reduced in ACRDYS2 fibroblasts. Based on the available data and experimental approaches, it is not possible to definitively exclude changes in the intrinsic activity of the mutant PDE4D protein that could contribute to the observed signalling differences. However, when total PDE4 activity is normalized to PDE4D protein levels (not shown), which are reduced in patient fibroblasts, the resulting values are comparable between patient and control cells. These findings are consistent with the reduction in total PDE4 activity being primarily attributable to decreased PDE4D expression, with compensatory activity from other PDE4 isoforms.

Basal cAMP levels appear to be similar in our control and patient fibroblasts, but were higher in ACRDYS2 fibroblasts without rolipram inhibition. This is consistent with the reduction in PDE4D levels of expression and consequently reduced enzymatic activity with increased cAMP / pCREB, that we have observed. However, it is not in line with the current view on the effect on PDE4D mutation in patients, where an overall loss of cAMP-PKA signalling is believed to occur. Kaname et al. reported reduced CREB phosphorylation in response to forskolin-induced synthesis of cAMP in ACRDYS2 EBV-immortalized patient lymphocytes as compared to controls, implying a reduced capacity for cAMP/PKA signalling in these cells [11]. In contrast, in our unmodified and unstimulated patient fibroblasts, pCREB was increased. Notwithstanding the different experimental settings and approach, it is notable that both the Kaname’s paper, the only other study where patient cells were used, though transformed, and our study support compensatory mechanisms as the molecular basis of ACRDYS2.

The finding of increased pCREB levels in ACRDYS2 fibroblasts raises several hypotheses that might explain CREB regulation in ACRDYS2 that should be explored in future studies. One possibility is that altered cAMP signalling may reduce CREB turnover rather than its activation state, as the pCREB/CREB ratio is unchanged, and there is no evidence of CREB regulation at the transcriptional level in ACRDYS2 fibroblasts as mRNA levels were not different between control and patient cells. In addition, the contribution of cAMP-responsive pathways other than PKA, such as EPAC signalling, to the regulation of CREB phosphorylation and activity in ACRDYS2 remains to be determined. Further studies will be required to establish the mechanisms underlying CREB dysregulation and its contribution to disease pathogenesis.

## Conclusions

In summary, this study provides the first analysis of endogenous PDE4D dysfunction in unmodified ACRDYS2 patient cells and reveals a previously unrecognized reduction in PDE4D expression at both the transcript and protein levels. The disease is associated with selective downregulation of key PDE4D isoforms, reduced PDE4 activity, increased pCREB, and compensatory upregulation of PDE4B, pointing to a complex cAMP signalling modulation. Our findings challenge the prevailing view that ACRDYS2 results solely from altered PDE4D catalytic function and instead suggest that transcriptional dysregulation and compensatory signalling responses are central features of the disease. Together, our results establish an important framework for understanding ACRDYS2 pathogenesis and identify mechanistic questions which will deserve future investigation.

## List of abbreviations

ACRDYS: Acrodysostosis
ACRDYS2: Acrodysostosis type 2
cAMP: cyclic adenosine monophosphate
CREB: cAMP responsive element binding protein
EBV: Epstein-Barr virus
EPAC: exchange protein directly activated by cAMP
iPPSDs: inactivating PTH/PTHrP signalling disorders
pCREB: phosphorylated CREB
PDE: phosphodiesterase
PDE4: phosphodiesterase 4
PDE4D: phosphodiesterase 4D
PKA: protein kinase A
PRKAR1A: type 1 regulatory subunit alpha of protein kinase A)
PTH: Parathyroid hormone
PTHrP: parathyroid hormone-related protein
UCR: upstream conserved region

## Acknowledgements

We would like to acknowledge valuable advice on imaging from Dr Dale Moulding, access to a human cell line from Prof. WL Di, and contribution to preliminary experiments from Tianshu Bai, Myriam Bahri and Valeriia Davydenko.

## Availability of data and materials

Data included in the publication can be made available on request.

## Authors contributions

OG generated the cells, characterized them, performed molecular characterization and wrote the manuscript; JL performed molecular characterization; EK performed immunofluorescence experiments; TM performed immunofluorescence experiments and some molecular characterization; LM and HL provided clinical insights and consent from patients for the use of biopsies; GB provided expert advice, antibodies and critical reading of the manuscript; PF obtained funding, supervised the work and wrote the manuscript.

## Supplementary Figure

**Supplementary Figure 1.**
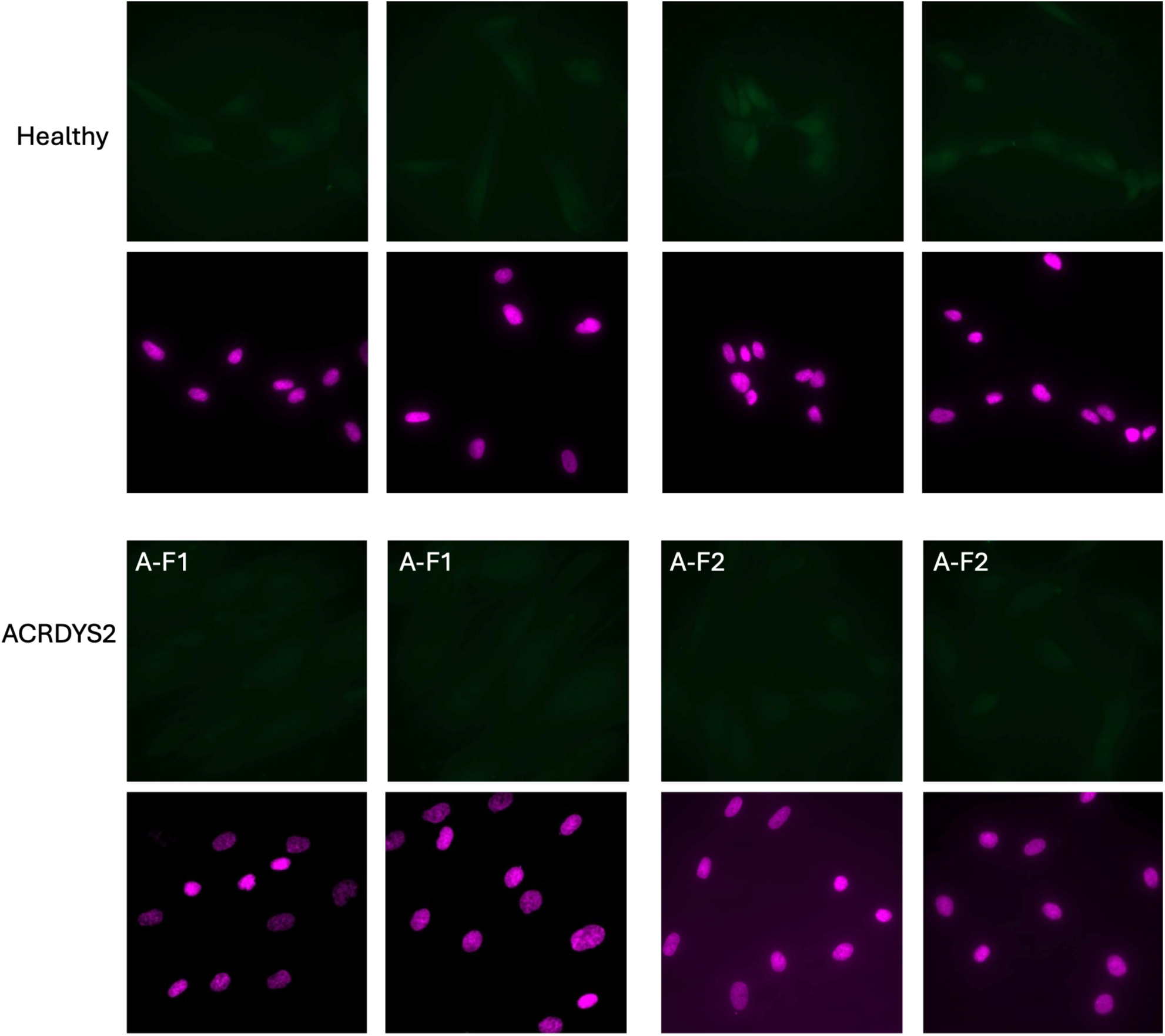
Negative controls for PDE4D immunostaining in healthy and ACRDYS2 fibroblasts. No staining is observed when the primary antibody to PDE4D is omitted. All images are at the same magnification; scale bar = 20 µm.

